# Metabolomic analysis of diverse mice reveals hepatic arginase-1 as source of plasma arginase in *Plasmodium chabaudi* infection

**DOI:** 10.1101/2021.06.11.448167

**Authors:** Nicole M. Davis, Michelle M. Lissner, Liliana M. Massis, Crystal L. Richards, Victoria Chevée, Avni S. Gupta, Frank C. Gherardini, Denise M. Monack, David S. Schneider

**Affiliations:** 1 Department of Microbiology and Immunology, Stanford University, Stanford, United States of America; 2 Laboratory of Bacteriology, National Institutes of Health, National Institute of Allergy and Infectious Diseases, Rocky Mountain Laboratories, Hamilton, Montana, United States of America

**Author notes:** Address correspondence to David Schneider.

**Keywords:** Malaria, Plasmodium chabaudi, Metabolomics, disease severity, arginine

## Abstract

Infections disrupt host metabolism, but the factors that dictate the nature and magnitude of metabolic change are incompletely characterized. To determine how host metabolism changes in relation to disease severity in murine malaria, we performed plasma metabolomics on eight *Plasmodium chabaudi*-infected mouse strains with diverse disease phenotypes. We identified plasma metabolic biomarkers for both the nature and severity of different malarial pathologies. A subset of metabolic changes, including plasma arginine depletion, match the plasma metabolomes of human malaria patients, suggesting new connections between pathology and metabolism in human malaria. In our malarial mice, liver damage, which releases hepatic arginase-1 (Arg1) into circulation, correlated with plasma arginine depletion. We confirmed that hepatic Arg1 was the primary source of increased plasma arginase activity in our model, which motivates further investigation of liver damage in human malaria patients. More broadly, our approach shows how leveraging phenotypic diversity can identify and validate relationships between metabolism and the pathophysiology of infectious disease.

**Importance:** Malaria is a severe and sometimes fatal infectious disease endemic to tropical and subtropical regions. Effective vaccines against malaria-causing *Plasmodium* parasites remain elusive, and malaria treatments often fail to prevent severe disease. Small molecules that target host metabolism have recently emerged as candidates for therapeutics in malaria and other diseases. However, our limited understanding of how metabolites affect pathophysiology limits our ability to develop new metabolite therapies. By providing a rich dataset of metabolite-pathology correlations, and by validating one of those correlations, our work is an important step toward harnessing metabolism to mitigate disease. Specifically, we showed that liver damage in *P. chabaudi*-infected mice releases hepatic arginase-1 into circulation, where it may deplete plasma arginine, a candidate malaria therapeutic that mitigates vascular stress. Our data suggest that liver damage may confound efforts to increase levels of arginine in human malaria patients.

## Introduction

Malaria is a serious infectious disease caused by Apicomplexan parasites in the genus *Plasmodium.* The most virulent species in humans is *P. falciparum*, which killed 405,000 people in 2018 (WHO). Malarial pathologies range from uncomplicated fever and anemia to severe and sometimes fatal conditions including severe anemia, metabolic acidosis, acute kidney injury, multi-organ failure, respiratory distress, and cerebral malaria (Stephens *et al*. 2012, Miller *et al*. 2013, Mackintosh *et al*. 2004, Leopold *et al*. 2019a, Leopold *et al*. 2019b).

Host metabolic changes frequently accompany malaria, often in association with severe systemic disease or organ-specific pathologies. For example, hepatocellular injury leads to elevated plasma levels of the metabolic enzymes aspartate and alanine transaminase (AST and ALT) (Ikemoto *et al*. 2001, Reuling *et al*. 2018). Kidney dysfunction also leads to plasma metabolic changes, the most characteristic being elevated plasma urea and creatinine (Hosten 1990, Conroy *et al*. 2016). Dysfunction in either organ can lead to organ failure and other systemic complications (Reuling *et al*. 2018). Hypoglycemia (White *et al*. 1983), metabolic acidosis (Leopold *et al*. 2019b), and other metabolic changes also accompany severe disease. However, it remains largely unclear how metabolic changes relate to the nature and degree of malaria pathophysiology.

Metabolites have recently emerged as important controllers of infection pathophysiology. Manipulation of glycolysis, for example, alters disease severity in malaria (Cumnock *et al*. 2018, Wang *et al*. 2018) and controls tissue damage in other infections (Wang *et al*. 2016). Other metabolic targets, including iron (Sanchez *et al*. 2018), immunomodulatory metabolic enzymes like arginase (Elahi *et al*. 2013, El Kasmi *et al*. 2008, Pesce *et al*. 2009), and metabolic hormones (Luan *et al*. 2019) have high potential to reduce pathology during infection. Undoubtedly, many metabolic therapies have yet to be discovered.

To uncover metabolic processes that are important during *Plasmodium* infection, we recently identified hundreds of plasma metabolites that change significantly when C57BL/6 mice are infected with *P. chabaudi* (Lissner *et al*. 2020). Many of these also change in human malaria (Gupta *et al*. 2017, Surowiec *et al*. 2015, Leopold *et al*. 2019a, Leopold *et al*. 2019b, Cordy *et al*. 2019, Lissner *et al*. 2020), but the causes and consequences of many of these metabolic changes remain unknown.

In a typical experiment, the field uses inbred strains to limit signal to noise to increase our probability of finding significant results. Here we used a collection of diverse inbred strains to deliberately introduce variation and then measure that variation. We selected eight inbred mouse lines (C57BL/6, WSB/EiJ, NZO/HILtJ, 129S1/SvImJ, A/J, CAST/EiJ, PWK/PhJ, and NOD/ShiLtJ) that represent 90% of laboratory mouse genetic diversity (Roberts *et al*. 2007) and vary widely in *P. chabaudi* infection severity as measured by survival, anemia, parasite load, temperature loss, and weight loss (**Fig. 1**). These Founder Strains were used to establish the Collaborative Cross (CC) and Diversity Outbred (DO) mouse populations (Churchill *et al*. 2012), which more closely approximate human genetic and phenotypic diversity than individual inbred lines. Collectively, these groups of diverse mice have improved our understanding of genetic and immune factors that alter the severity of infections like influenza and tuberculosis (Leist *et al*. 2016, Elbahesh & Schughart 2016, Noll *et al*. 2019, McHugh *et al*. 2013, Niazi *et al*. 2015, Kurtz *et al*. 2020). Our goal was to use diverse mice to identify metabolic factors that alter infection severity in malaria.

**Figure 1.**
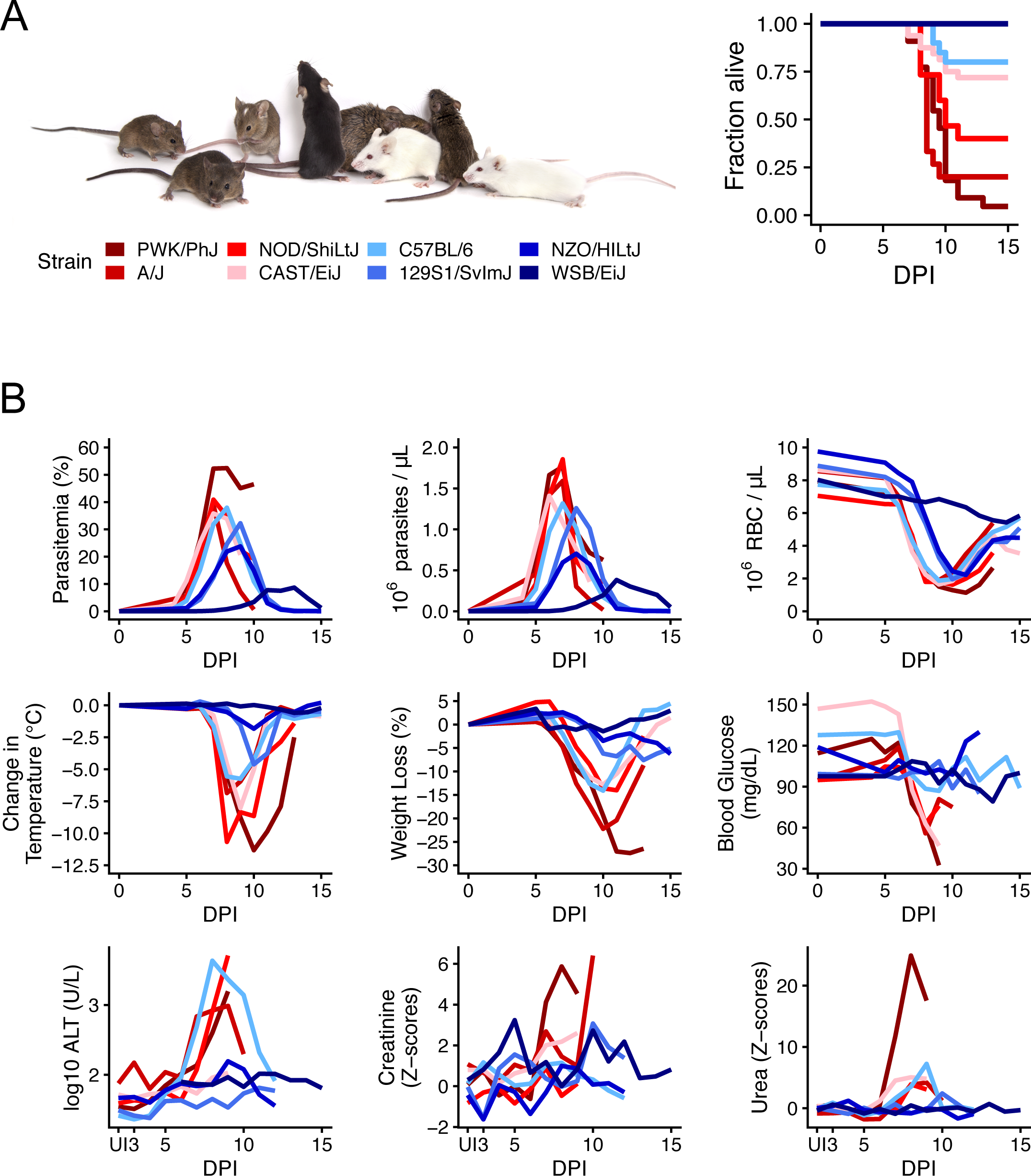
Malarial disease severity varies widely across eight genetically diverse mouse strains. **(A)** The 8 Founder Strains (left to right): CAST/EiJ, 129S1/SvImJ, WSB/EiJ, C57BL/6J, NZO/HILtJ, NOD/ShiLtJ, PWK/PhJ, A/J. Survival: Strain colors reflect average survival and parasite load, with dark blue the most resilient/fewest parasites, and dark red the least resilient. (n=10 for 100% resilient WSB/EiJ, NZO/HILtJ, 129S1/SvImJ; n=30 for 73% resilient C57BL/6, n=32 for 72% resilient CAST/EiJ, n=15 for 40% resilient NOD/ShiLtJ, n=15 for 20% resilient A/J, and n=22 for 4.5% resilient PWK/PhJ). DPI = days post-infection. Photo reproduced with permission from Jackson Laboratories. **(B)** Nine metrics of disease severity across the eight Founder Strains over the course of acute *P. chabaudi* infection. Parasitemia = % parasitized RBCs, parasite density = 10^6^ RBCs per microliter blood, anemia = 10^6^ RBCs per microliter blood). Lines indicate the median of >= 5 mice per strain per day. Z-scores = standard deviations from the mean of uninfected C57BL/6 mice.

To link malaria pathophysiology with host metabolic changes, we measured plasma metabolites and markers of pathology and immune responses in the eight *P. chabaudi*-infected Founder Strains. Using principal component analysis, we found that the magnitude of metabolic response to *P. chabaudi* infection was directly related to disease severity. Using correlation analysis, we found that arginine depletion was a specific marker for malaria-induced liver damage. We identified additional candidate molecules for future study. We showed that *P. chabaudi*-induced liver damage releases arginine-consuming hepatic arginase-1 into circulation, which explains increased plasma arginase activity in our model and may explain elevated plasma arginase in other infections. Our results support the use of diverse mice to understand the links between metabolism and disease severity during infection, and motivate further investigation of the metabolic consequences of liver damage in human malaria.

## Results

### Infection alters metabolomic profiles in diverse mice and humans

We previously identified 370 metabolites that change significantly during *P. chabaudi* infection in C57BL/6 mice (Lissner *et al*. 2020). However, wildtype C57BL/6 mice are largely resilient to infection, which limited our ability to understand how host metabolism changes in severe disease. To measure metabolism in mice with a broader range of disease severities, we used the eight Founder Strains (C57BL/6, WSB/EiJ, NZO/HILtJ, 129S1/SvImJ, A/J, CAST/EiJ, PWK/PhJ, and NOD/ShiLtJ), which vary widely in survival following *P. chabaudi* infection (**Fig. 1a**). For each strain, we monitored disease severity, plasma metabolites, plasma cytokines, and circulating immune cells during acute infection (days 5-12 post-infection or days 5-17 for WSB/EiJ mice), when metabolic perturbations in C57BL/6 mice are greatest (Lissner *et al*. 2020). The Founder Strains varied widely across nine metrics of malaria severity that largely tracked with survival and parasite load: anemia (red blood cells (RBCs) per microliter of blood), hypothermia, weight loss, hypoglycemia, liver injury (plasma ALT), and kidney injury (plasma urea, creatinine) (**Fig. 1b**). Plasma cytokines tended to be higher in resilient strains than in non-resilient strains, with two exceptions: cytokines in resilient WSB/EiJ mice remained low throughout infection (**Fig. S1a**), and two non-resilient strains (PWK/PhJ and NOD/ShiLtJ) mounted hyperinflammatory responses (e.g. high IFN-*γ*, TNF-*α*, IL-6, and low IL-10, TGF-*β*) on days 8-9. Immune cell responses were largely consistent across strains, with elevated T cells, B cells, NK cells, and monocytes between days 7-9 post-infection (**Fig. S1b**). NK cells were particularly high in resilient NZO/HILtJ mice, as were T and B cells in resilient 129S1/SvImJ and non-resilient CAST/EiJ mice, respectively. Collectively, our data highlight the diversity of disease and immune phenotypes in the eight Founder Strains.

Inter-strain variation in malarial disease severity allowed us to ask several questions about the relationship between disease severity and metabolism: is overall metabolic disruption greater in sicker animals? How do individual metabolites relate to different facets of disease? Can these metabolic markers of pathology provide new mechanistic insights into the pathophysiology of malaria or other infectious diseases?

To answer how disease severity impacted the overall composition of plasma metabolomes, we performed dimensionality reduction and visualization of 635 metabolites (see Methods) using principal component analysis (PCA). Variance in PC2 (11.2%) was associated with baseline differences among mouse strains, with NZO/HILtJ samples having the highest PC2 values and wild-derived strains (CAST/EiJ, WSB/EiJ, and PWK/PhJ) having the lowest PC2 values (**Fig. 2a**). Variance in PC1 (31.2 %) was associated with infection severity. Highly resilient mice (WSB/EiJ, NZO/HILtJ, and 129S1/SvImJ) had high PC1 values throughout infection, while non-resilient strains had low PC1 values during periods of acute illness (days 7-10). Phase curves of PC1 versus parasite density show larger loops in disease space for less resilient strains, as is expected for variables that influence disease severity (**Fig. 2b**, Torres *et al*. 2016). We next used canonical correspondence analysis (CCA), a technique similar to PCA, to identify molecules that drove the most variation among samples (**Table S1**). Molecules that differentiated sick from healthy samples were associated with altered feeding behavior (food components, ketosis markers, indoles, serotonin, acylcarnitines), oxidative stress (glutathione, 2-hydroxyisobutyrate), and liver and kidney dysfunction (formiminoglutamate, bilirubin, homocitrulline, urea, p-cresol glucuronide, p-cresol sulfate, N-acetyltyrosine). We also performed Student’s t-tests to identify molecules that differentiate resilient and non-resilient mice before infection (**Table S2**). Plasma levels of a number of metabolites differed significantly between uninfected resilient and non-resilient mice, including the food component and osmolyte betaine (Lever & Slow 2010), sphingomyelins, lysine, and pantothenic acid.

**Figure 2.**
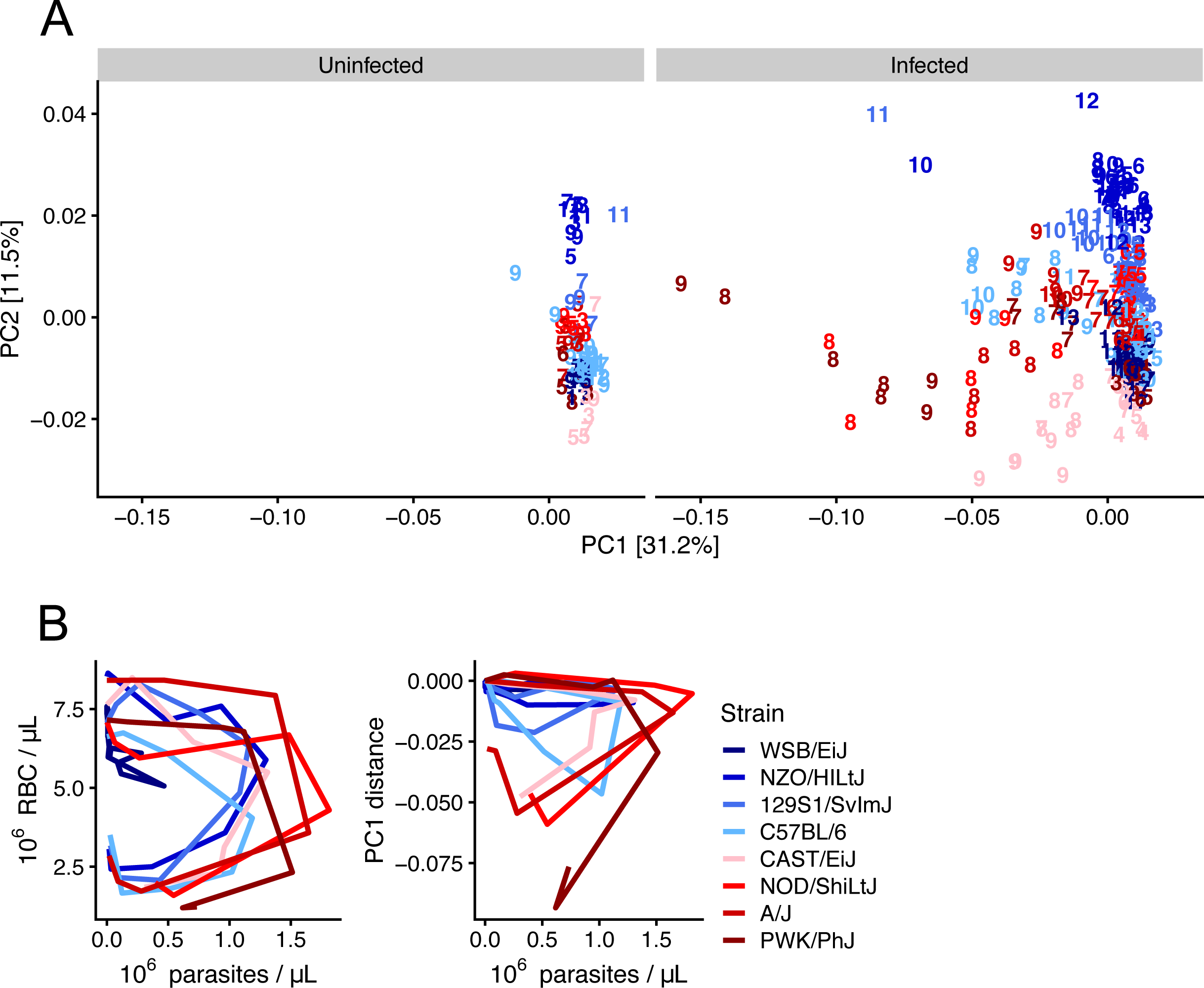
Genetic background and *Plasmodium* infection influence murine plasma metabolomes. **(A)** Bray-Curtis dissimilarity index was computed on 635 metabolites per sample for infected and uninfected samples. Principal component analysis (PCA) was performed, and sample positions in the first two principal components are displayed. PC1 and PC2 explain 31.2% and 11.5% variation, respectively. Each mouse sample is represented as a number that indicates days post-infection (or mock infection). **(B)** Disease space loops of RBCs and median PC1 values (derived from Fig. 2 data) plotted against median parasite densities throughout acute *P. chabaudi* infection. Each line represents the median value for >= 5 mice per strain per day (except final days for non-resilient strains in which <5 individuals remain).

### Correlations identify metabolic markers for disease severity in *P. chabaudi* malaria

Having determined that the sickest mice displayed the greatest changes in plasma metabolome composition, we next obtained a more granular understanding of pathology-metabolism relationships by correlating individual metabolites with our nine metrics of disease (**Fig. 3a**). The kidney dysfunction marker urea had the highest number of strong correlations (R^2^ > 0.4), followed by ALT, creatinine, and temperature loss. This is consistent with results from our PCA, which identified metabolic markers for organ dysfunction as strong drivers of variation in PC1. Anemia and hypoglycemia also correlated well with some plasma metabolites, while few metabolites correlated well with weight loss, parasitemia, or parasite density. Altogether, our correlations suggest the host is a stronger driver of infection-induced plasma metabolic changes than the parasite.

**Figure 3.**
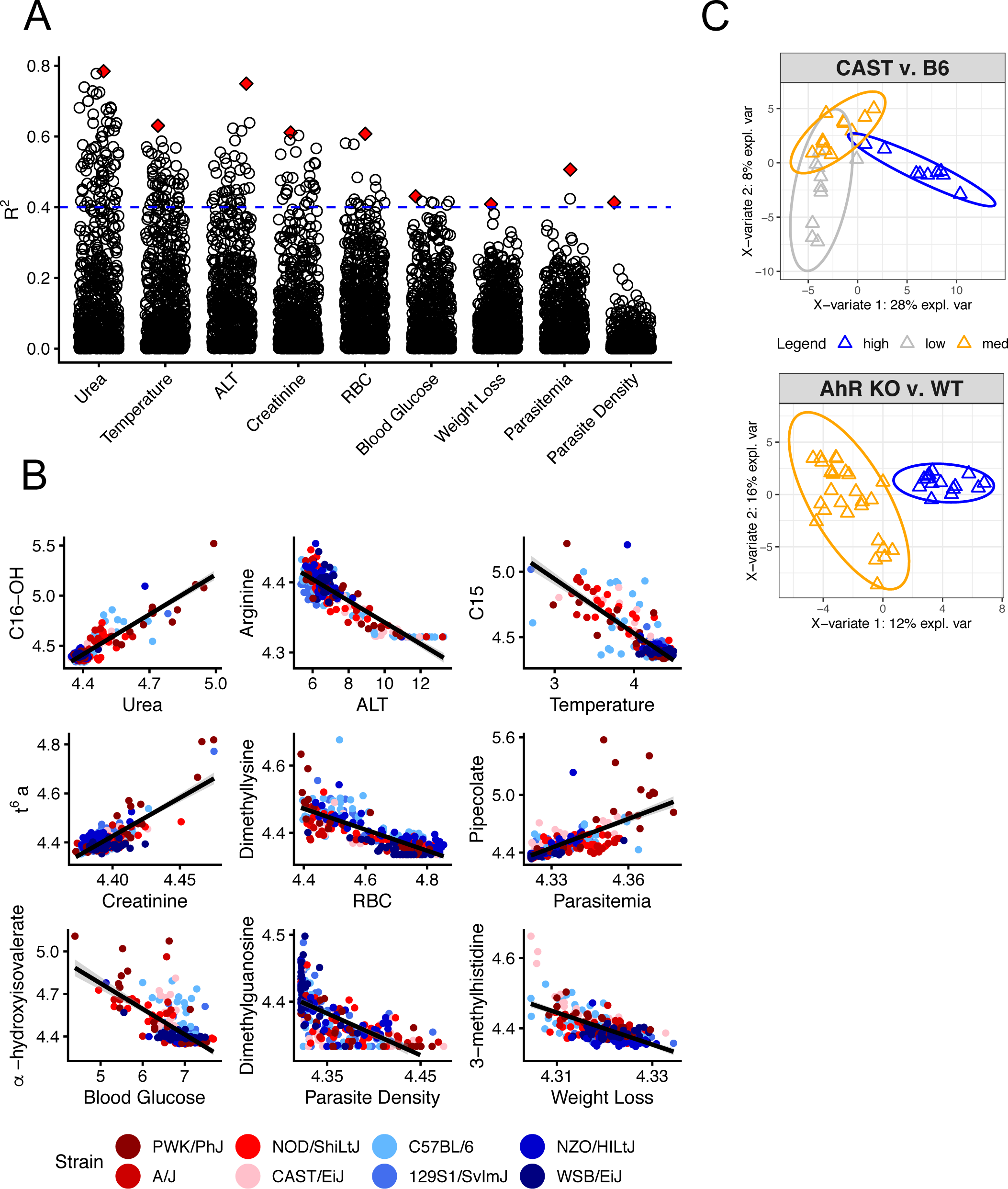
Metabolic indicators of disease severity in P. chabaudi-infected mice. **(A)** Log2-transformed scaled imputed ion counts (n=751 metabolites) were correlated with the nine metrics of disease severity shown in Figure 1. Each point represents the R^2^ value of the Pearson correlation for each metabolite and health combination (n= 751 x 9 total correlations). Red diamonds indicate the best correlation for each health metric. The dotted blue line is the threshold for R^2^ = 0.4. **(B)** Correlation plots with best-fit lines for red diamonds in A. C16-OH = hydroxypalmitoylcarnitine, C15 = pentadecanoylcarnitine, TMAP = N, N, N-trimethyl-L-alanyl-L-proline betaine. R^2^ values (left to right): (top row) 0.78, 0.63, 0.75, (middle row) 0.61, 0.61, 0.43, (bottom row) 0.41, 0.51, 0.41. **(C)** sPLS-DA separates samples (triangles) from *P. chabaudi*-infected C57BL/6 and CAST/EiJ mice on the basis of liver damage (blue = high, orange = moderate, gray = low). A similar analysis separates samples from *P. chabaudi*-infected *Ahr^+/+^* and *Ahr^-/-^* C57BL/6 mice on the basis of liver damage (high and moderate liver damage, respectively).

Some metabolites correlated well with multiple disease metrics. For example, *α*-hydroxyisovalerate, a hallmark of defective leucine metabolism (Tanaka *et al*. 1966) and the best indicator of hypoglycemia in our malarial mice (**Fig. 3b**), correlated well with all disease metrics except parasite load (**Dataset S2**). Similarly, elevated long-chain acylcarnitines were the best predictors of high plasma urea levels and temperature loss (**Fig. 3b**), but many also positively correlated with ALT. Long-chain acylcarnitines are also elevated in other pathological states like LPS-induced inflammation (Ganeshan *et al*. 2019) and acute kidney injury (Andrianova *et al*. 2020) (**Fig. S2**). Levels of *α*-hydroxyisovalerate and long-chain acylcarnitines also increase during starvation (Pietrocola *et al*. 2017, Holecek 2018, **Fig. S2**), which is consistent with our previous observations of *P. chabaudi*-induced anorexia (Cumnock *et al*. 2018). Collectively, these multi-correlative metabolites may point to shared underlying mechanisms of metabolic disruption across disease states.

In contrast to metabolites that were broadly indicative of severe disease, some metabolites specifically correlated with just one pathology. Some of these associations have been identified previously, including pipecolate as a marker for parasitemia (Beri *et al*. 2019) and N,N,N-trimethyl-alanylproline betaine, which positively correlates with creatinine in our data and was recently identified as a sensitive marker for kidney injury (Velenosi *et al*. 2019). Other metabolites suggested directions for future study. For example, 3-methylhistidine, a proposed marker for muscle protein breakdown (Elia *et al*. 1981), correlated with weight loss in our malarial mice. Muscle protein breakdown has not been reported in *P. chabaudi*-infected mice, but cachexia occurs in other inflammatory conditions (Schieber *et al*. 2015, Luan *et al*. 2019, Melchor *et al*. 2020). We also noted a strong association between arginine depletion and elevated ALT (**Fig. 3b**), a molecular proxy for liver damage (Ikemoto *et al*. 2001, Reuling *et al*. 2018). While both abnormalities are prevalent in human malaria and pose health concerns (Yeo *et al*. 2007, Yeo *et al*. 2009, Weinberg *et al*. 2016, Gupta *et al*. 2017, McDonald *et al*. 2018, Reuling *et al*. 2018), the connection between liver damage and arginine in malaria is understudied.

### Arginine depletion is a specific indicator of liver damage in *P. chabaudi* malaria

To better determine how specifically plasma arginine depletion indicates liver damage, we applied sparse partial least squares discriminant analysis (sPLS-DA) to identify the best metabolic markers for high ALT in our data and in a second dataset (Lissner *et al*. 2020). sPLS-DA is a supervised learning method that selects variables that best differentiate samples based on user-assigned groups, and is well-suited to sparse and heterogeneous multi-omics data (Le Cao *et al*. 2011, Rohart *et al*. 2017, Fletcher *et al*. 2018, **Methods**). We limited analysis in our dataset to two strains of mice – CAST/EiJ and C57BL/6 – that had low and high levels of ALT, respectively, but similar levels of other disease severity metrics. By comparing samples from only these two strains, we reasoned we could identify metabolites that are specific to liver damage rather than other pathologies. Our sPLS-DA model identified arginine, adenosine, and methionine along with other metabolites in component 1 as best able to discriminate among samples with “high”, “medium”, or “low” ALT values (**Fig. 3c**, top, **Table S3**). We performed a similar analysis (**Fig. 3c**, bottom) on plasma metabolomes from *P. chabaudi*-infected wildtype and *Ahr^-/-^* C57BL/6 mice, which have high and low levels of ALT during infection, respectively. Upon comparing metabolic candidates from sPLS-DA and our correlation analysis (n=40-50 metabolites for each of three analyses), we found that overlap between the three analyses was low – just 1-3 metabolites were identified by any two of three analyses – but arginine was selected by all three (**Fig. 3c**, **Dataset S2, Table S3**).

### Malarial liver damage releases hepatic Arg1 into circulation in conjunction with arginine depletion

In healthy, ureotelic animals, hepatic arginase-1 (Arg1) converts arginine to ornithine and urea (Morris 2016, **Fig. 4a**) in a process that is spatially restricted to hepatocytes. However, drug-induced hepatocellular injury releases hepatic arginase-1 into circulation where it depletes plasma arginine (Ikemoto *et al*. 2001). Only associative evidence has linked arginine depletion with hepatic arginase and liver injury in malaria (Yeo *et al*. 2009). Given the strong and specific association between arginine depletion and ALT in our data, we hypothesized that *P. chabaudi*-induced liver damage depletes plasma arginine by releasing hepatic Arg1 into circulation. In support of our hypothesis, we found that the arginase product ornithine is elevated in mice with high ALT and low arginine (**Fig. 4b**, top row: C57BL/6, A/J, NOD/ShiLtJ, PWK/PhJ). In contrast, the urea cycle metabolite and nitric oxide synthase product citrulline varied independently of arginine depletion and ALT. Further analysis of Founder Strain samples revealed a positive correlation between plasma arginase activity and plasma ALT (R^2^=0.75, **Fig. 4c**).

**Figure 4.**
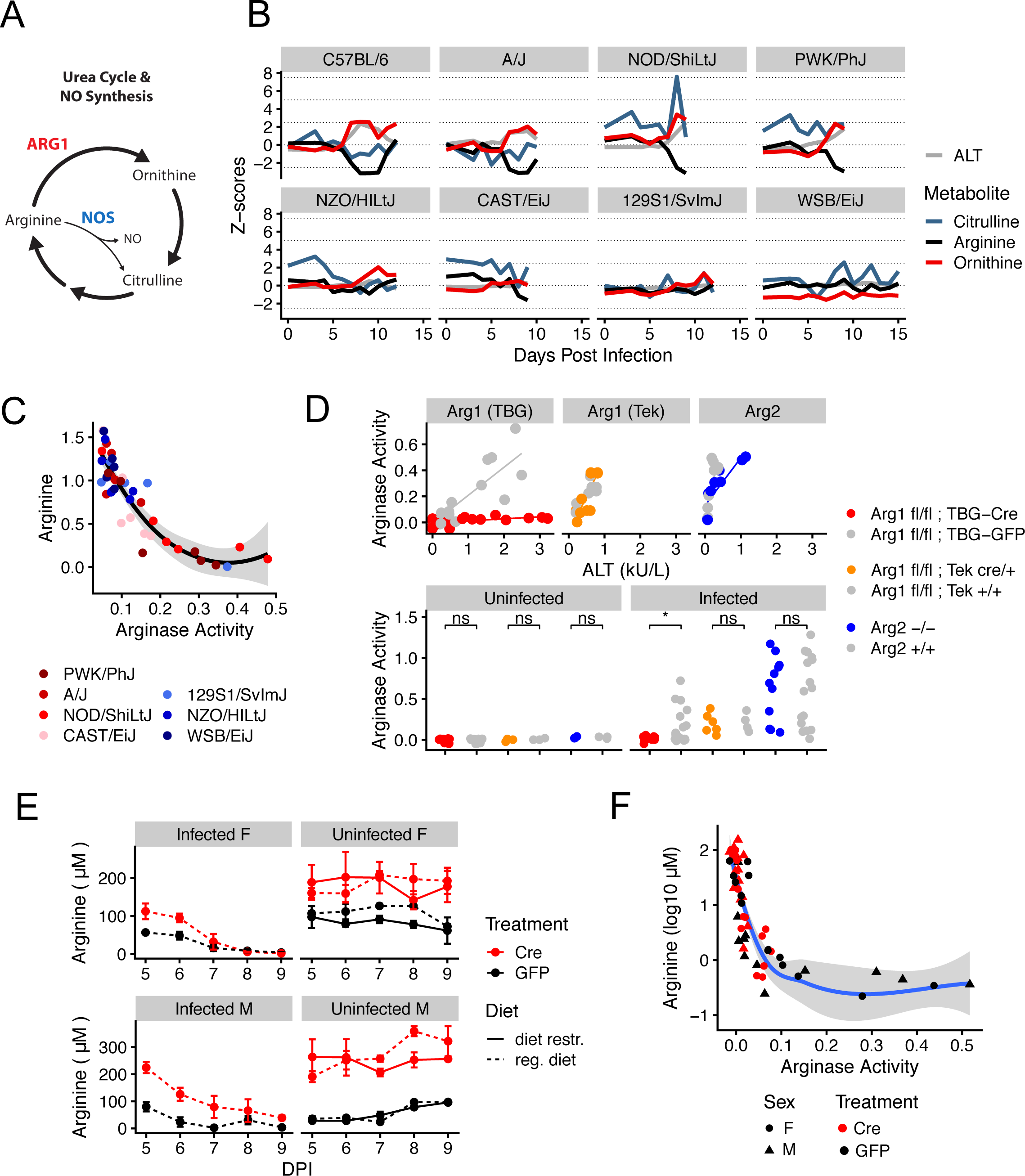
The liver controls arginine metabolism via arginase-1. **(A)**A schematic of the urea cycle, polyamine metabolism, and nitric oxide production. NO = nitric oxide, NOS = nitric oxide synthase, ARG1 = arginase-1. **(B)** Median scaled imputed ion counts of arginine, ornithine, and citrulline throughout infection. Z-score = 1 standard deviation from the mean of uninfected C57BL/6 mice. n=5 animals per strain per day. **(C)** Plasma arginase activity (nmol H_2_O_2_ / min. / uL blood) and scaled imputed ion counts of arginine (log10-transformed Z-scores) in 7 of 8 *P. chabaudi*-infected Founder Strains (n=6 per strain). Loess curve with standard error. **(D)** Plasma arginase activity (nmol H_2_O_2_ / min. / uL blood) and ALT in arginase knockout C57BL/6 mice before and during *P. chabaudi* infection (n>=3 uninfected mice per genotype, n>=6 infected mice per genotype in 2 experimental replicates). Welch’s t-test: *p<0.05, **p<0.01. **(E)** Mean (+/- SE) plasma arginine in uninfected (+/- dietary restriction) and *P. chabaudi*-infected *Arg1^fl/fl^; TBG-Cre* and *Arg1^fl/fl^; TBG-GFP* mice (n=3 per condition per sex, day 5 post-infection is day 12 post-AAV injection). **(F)** Plasma arginase activity (nmol H_2_O_2_ / min. / uL blood) and arginine for infected mice and days shown in E. Loess curve with standard error.

Plasma arginase activity is increased in malaria patients, but it has been attributed to other sources of Arg1 such as red blood cells or monocytes (Yeo *et al*. 2009, Weinberg *et al*. 2016). The other mammalian arginase arginase-2 (Arg2), which is highly expressed in immune cells as well as the gut and kidney (Shi *et al*. 2001, Fritz 2013, Lindez *et al*. 2019), is rarely considered (Weinberg *et al*. 2016). To determine the isoform and tissue source of plasma arginase during *P. chabaudi* infection, we utilized a globally Arg2-deficient and two tissue-specific Arg1-deficient C57BL/6J mouse strains. Because murine Arg1 deletion results in lethality 14 days after birth (Iyer *et al*. 2002), we crossed Arg1 floxed mice (*Arg1^fl/fl^* mice, El Kasmi *et al*. 2008) with *Tek-Cre* mice, generating *Arg1^fl/fl^; Tek-Cre* mice that lack Arg1 specifically in blood cells and the endothelium. To overcome lethality caused by liver-specific Arg1 deficiency (Ballantyne *et al*. 2016), we injected *Arg1^fl/fl^* mice with adeno-associated viral (AAV) vectors that express Cre (or GFP for control animals) under control of the liver-specific *TBG* promoter. This yielded *Arg1^fl/fl^; TBG-Cre* mice that display hyperargininemia and reduced plasma arginase activity at 2 weeks post-injection, and that die from liver Arg1 deficiency at 3 weeks post-injection (**Fig. S3a,b**).

We infected these knockout mice and their respective controls with *P. chabaudi* and measured ALT and plasma arginase activity at peak infection intensity. As expected, uninfected mice of all genotypes exhibited low ALT and arginase activity. In infected, non-knockout mice, ALT and arginase activity positively correlated (**Fig. 4d**). ALT and arginase activity also correlated in *Arg2^-/-^* and *Arg1^fl/fl^; Tek-Cre* mice, suggesting neither Arg1 in the blood and endothelium nor Arg2 contribute to plasma arginase activity during *P. chabaudi* infection. In contrast, *Arg1^fl/fl^; TBG-Cre* mice exhibited high levels of ALT but significantly reduced plasma arginase activity relative to infected *Arg1^fl/fl^; TBG-GFP* mice (**Fig. 4**). This suggests hepatic Arg1 is the source of plasma arginase activity in *P. chabaudi*-infected mice.

Low plasma arginine is associated with increased vascular stress (Yeo *et al*. 2009) and negative pregnancy outcomes in malaria patients (McDonald *et al*. 2018). However, arginine infusion in malaria patients sometimes fails to restore arginine levels and mitigate vascular stress (Yeo *et al*. 2013), motivating further exploration of the causes of hypoargininemia. Plasma arginase is associated with arginine depletion in malaria (Weinberg *et al*. 2016, Yeo *et al*. 2009), which is consistent with our hypothesis that hepatic Arg1 depletes arginine. Infection-induced anorexia and altered flux of arginine through the urea cycle are other explanations that accounted for some but not all arginine depletion in mice with experimental cerebral malaria (Alkaitis *et al*. 2016). To assess the importance of these factors on plasma arginine dynamics, we restricted food intake in uninfected *Arg1^fl/fl^; TBG-Cre* and *Arg1^fl/fl^; TBG-GFP* mice to the level of age- and sex-matched *P. chabaudi*-infected mice (**Fig. S3c**). Despite comparable weight loss in infected and uninfected mice (**Fig. S3c**), decreases in plasma arginine during dietary restriction were relatively modest; only infected mice displayed dramatic hypoargininemia (**Fig. 4e**) Arginine depletion correlated strongly with plasma arginase activity in both genotypes, even in *Arg1^fl/fl^; TBG-Cre* mice with significantly reduced arginase activity (**Fig. 4f**), suggesting that even residual amounts of circulating arginase are sufficient to reduce plasma arginine by 100- fold or more. Finally, given the dramatic elevation of plasma arginine following knockdown of hepatic Arg1 (**Fig. S3a**, **Fig. 4e**, Ballantyne *et al*. 2015), it is unlikely that this type of urea cycle disruption contributes to arginine depletion. Collectively, our data suggest that hepatic Arg1 maintains plasma arginine homeostasis and that circulating hepatic Arg1 depletes plasma arginine during *P. chabaudi* infection.

### Metabolic changes in murine malaria are seen in other infections

Altered host metabolism is ubiquitous in infectious disease, which led us to ask if *P. chabaudi* infection shares metabolic features with other infections. We started by comparing our metabolic data to a similar metabolomics experiment performed on DO mice infected with *Salmonella enterica* serovar Typhimurium. Using PCA and CCA, we determined that infection altered plasma metabolomes in *S.* Typhimurium infection (**Fig. S4a**), but the metabolites that drove variation in the data did not overlap with the metabolites driving variation in murine malarial metabolomes. However, a subset of *Salmonella*-infected DO mice exhibited arginine depletion that correlated with elevated plasma arginase activity (**Fig. S4b**), suggesting that both *Plasmodium* and *Salmonella* infections result in arginase-mediated arginine depletion.

We next compared metabolic changes in *P. chabaudi* to human malaria metabolomes. We started with publicly available metabolic data from two populations of *P. falciparum*-infected individuals: Thai adults with uncomplicated malaria (MaHPIC Consortium) and Malawian pediatric patients with severe cerebral malaria (Gupta *et al*. 2017). Using metabolites that were measured in both human studies and our mouse study (predominantly amino acids and lipids), we created a network using the correlation matrix of mouse samples at peak disease severity (days 8-9) and all human samples (**Fig. 5a**). In the network, mice with mild disease were better connected to Thai adults with uncomplicated malaria, while mice with severe disease correlated better with Malawian pediatric patients with severe disease. A subset of mouse samples (C57BL/6 at peak infection severity) was highly connected to samples from both human datasets, suggesting they shared similarity with both human populations.

**Figure 5.**
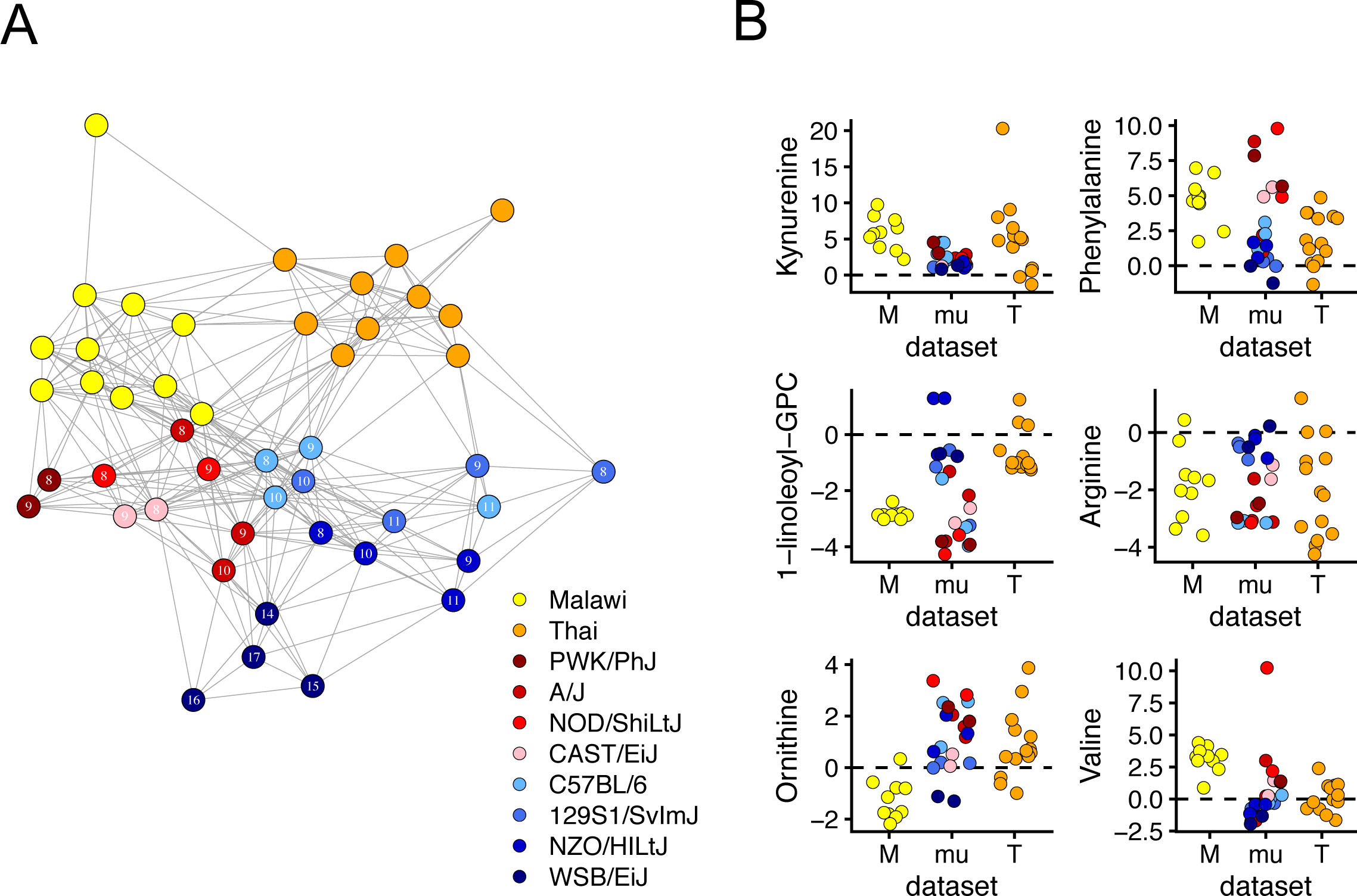
Mice and humans share plasma metabolic features of *Plasmodium* infection. A mouse-human correlation network highlights similarity of mouse and human metabolic responses to malaria. Two human metabolic datasets from Malawian cerebral pediatric *P. falciparum*-infected malaria patients (yellow, n=10) and adult, Thai severe *P. falciparum*-infected malaria patients (orange, n=10) were analyzed. **(A)** The 42 metabolites (values Z-scored from uninfected mice or patients) measured in all three datasets were included in a sample-sample inter-dataset correlation analysis. The resulting network is shown. Each sample is a network node, and edges were drawn between any sample with R^2^ >= 0.5. Mouse samples are the median values for each strain (day 8-11 or 14-17 for WSB/EiJ) post-infection. Each human sample represents one individual. **(B)** The same data are used to display for 6 individual metabolites.

To understand which metabolic changes drove inter-species similarity in the correlation network, we compared the levels of individual metabolites across our mice and the human samples (**Fig. 5b**). In most mice and humans, infection increased levels of phenylalanine and kynurenine and decreased levels of arginine, glycine, serine, and long-chain glycerophosphatidylcholines (GPCs). Notably, some metabolites like valine and 1-linoleoyl-GPC changed more in severe disease (Malawian patients and less resilient mouse strains, reds) than in mild disease (Thai patients and resilient mouse strains, blues). For many other metabolites, the two human datasets qualitatively differed while mice tended to either match one population or display intermediate responses (**Fig. 5b**). For example, ornithine was elevated in Thai patients and our mice, but depleted in Malawian patients. Conversely, valine was elevated in Malawian patients and mice, but relatively unchanged in Thai patients.

## Discussion

Host metabolism changes in response to infection, but we still know little about how host metabolism responds to severe versus mild disease. To better understand how disease severity affects host metabolism in murine malaria, we monitored disease severity metrics (**Fig. 1**) and plasma metabolites in eight genetically diverse mouse strains that vary widely in survival following acute *Plasmodium chabaudi* infection. While *P. chabaudi* primarily causes nonfatal severe anemia due to high parasitemia, we observed a wide range of other disease severity phenotypes in which parasitemia, anemia, temperature loss, weight loss, liver and kidney injury, and hypoglycemia were generally more severe in mice that succumb to infection. We also observed a wide range of metabolic responses, and used PCA to show that the magnitude of metabolic response scaled with overall disease severity (**Fig. 2**).

Previous human studies (e.g. Surowiec *et al*. 2015, Leopold *et al*. 2019a, Leopold *et al*. 2019b) also reported malaria severity-dependent changes in host metabolism. To our knowledge, Leopold *et al*. provide the most comprehensive picture of these changes in Bangladeshi patients, both in terms of the number of metabolites assessed and in number of facets of disease severity examined, including acute kidney injury, metabolic acidosis, and coma. Our work complements these studies by including additional metrics of disease severity with quantitative data for the degree of disease severity. Using our approach, we showed that malarial mice and humans share some plasma metabolic changes during infection (**Fig. 5**). These conserved metabolic changes fell into one of two broad categories: responses consistent in all three populations, and those shared by mice and just one of the two human populations.

All hosts shared some malaria-induced metabolic changes. Kynurenine and phenylalanine were elevated in all individuals, mouse and human. Kynurenine is a tryptophan breakdown product and ligand of the aryl hydrocarbon receptor (AhR), which protects the host in multiple models of malaria and sepsis (Lissner *et al*. 2020, Brant *et al*. 2014, Bessede *et al*. 2014). Phenylalanine is also elevated during infection, as in other malaria studies (Enwonwu *et al*. 1999, Lopansri *et al*. 2006, Leopold *et al*. 2019a). Phenylalanine elevation may be caused by increased protein catabolism (Alkaitis *et al*. 2016) and malaria-induced depletion of tetrahydrobiopterin (Yeo *et al*. 2015), a cofactor required for enzymatic conversion of phenylalanine to tyrosine (Kure *et al*. 2004). Arginine was depleted in most individuals, which is consistent with all mouse and human studies we examined (Yeo *et al*. 2007, Yeo *et al*. 2009, Surowiec *et al*. 2015, Alkaitis *et al*. 2016, Gupta *et al*. 2017, Leopold *et al*. 2019a, Leopold *et al*. 2019b, Rubach *et al*. 2019). Finally, most infected mice and humans had depleted long-chain GPCs (also observed in Surowiec *et al*. 2015, Cordy *et al*. 2019), which may reflect changes in cell membranes, cell membrane lysis, or altered lipid metabolism. The non-essential amino acids glycine and serine, which *P. falciparum* uses in folate derivative-dependent DNA synthesis (Nzila *et al*. 2005), were also depleted. Given the similarities we identified between humans and mice, our work supports the use of *P. chabaudi*-induced murine malaria as a model to understand host metabolic responses to malaria.

For a number of metabolites, mice and one human population differed from the second human population. For instance, ornithine was generally higher in our malarial mice and in Thai adults with uncomplicated malaria (MaHPIC), but low in Malawian children with cerebral malaria (Gupta *et al*. 2017). We anticipate elevated plasma arginase explains ornithine elevation in malarial humans as it does in our murine data. Like ornithine, long-chain acylcarnitines (ACs) (e.g. C16 and C18) were also elevated in malarial mice and Thai patients, but depleted in Malawian patients (data not shown). We found that long-chain ACs correlated with many pathologies in our model, including liver and kidney dysfunction and temperature loss. Others have also noted elevation of long-chain ACs following acute kidney injury, fasting, and LPS injection in mice (**Fig. S2a**) (Andrianova *et al*. 2020, Pietrocola *et al*. 2017, Ganeshan *et al*. 2019). Fasting elevates long-chain ACs and is usually accompanied by an increase in fatty acid *β*-oxidation gene expression (Pietrocola *et al*. 2017). However, in infection (Ganeshan *et al*. 2019) and long-chain fatty acid oxidation disorders (Knottnerus *et al*. 2018), *β*-oxidation is suppressed. Measuring fatty acid *β*-oxidation gene expression in *P. chabaudi*-infected mice would help determine whether elevated long-chain ACs in our model indicate a normal fasting response or impaired *β*-oxidation. We also found that the branched-chain amino acids (BCAAs) leucine, isoleucine, and valine were elevated in mice with severe disease. Leopold *et al*. (2019a) also noted BCAA elevation in patients with severe but not uncomplicated malaria, which is consistent with our mice and the two human datasets examined here. BCAAs are altered in fasting states in both mice and humans (Pietrocola *et al*. 2017). Though not present in either human dataset, *α*-hydroxyisovalerate was a sensitive marker for hypoglycemia and weight loss in our murine data, and it is also indicative of defective leucine metabolism (Tanaka *et al*. 1966). Collectively, these data suggest infection-induced anorexia or changes in energy metabolism produce a strong molecular signature in severe malaria.

Our data suggest that diverse mice capture more of the diversity of human malaria responses than one human population alone can. However, the small number of metabolites and datasets (3) included in this study limit our ability to make broad generalizations. We also note that differences in age, geography, study design, and other variables can introduce confounding factors that affect metabolism in the two human populations examined here – Malawian children and Thai adults. We selected these datasets because they had the highest number of metabolites available for comparison with our murine data. Future studies would benefit from measuring samples from multiple mouse and human populations in tandem.

In addition to metabolites that generally indicate severe disease, several molecules from our study associated with individual pathologies (**Fig. 3**). Two of these correlations corroborated known associations: pipecolate as a biomarker for parasitemia (Beri *et al*. 2019), and N,N,N-trimethyl-alanylproline betaine (TMAP), which correlated with the kidney injury marker creatinine and was recently identified as a sensitive marker of kidney dysfunction (Velenosi *et al*. 2019). Other correlations like the association of 3-methylhistidine with weight loss and the correlation between *α*-hydroxyisovalerate and hypoglycemia are consistent with previous reports of muscle protein catabolism elevating plasma 3-methylhistidine (Elia *et al*. 1981), and of defective leucine metabolism causing hypoglycemia and *α*-hydroxyisovalerate elevation (Tanaka *et al*. 1966). Together, these findings support the validity of our correlation method, and motivate further study of associations like dimethyllysine and red blood cells (**Fig. 3b**), *N*^6^-threonylcarbamoyladenosine (t^6^A) and creatinine (**Fig. 3b**), acylcarnitines (**Fig. 3b**) as discussed above, and others.

We demonstrated that our bioinformatic approach yielded precise and testable mechanistic hypotheses. Specifically, our correlation analysis pointed to the liver, rather than red blood cells, immune cells, or the parasite as the source of arginine-depleting arginase (**Fig. 3b-c**). We tested this hypothesis by measuring plasma arginase and arginine in liver-specific Arg1 knockout mice, blood and endothelial-specific Arg1 knockout mice, and Arg2 knockout mice on a C57BL/6 background. In support of our bioinformatic data, we found that plasma arginase activity in malaria was reduced only in liver-specific Arg1 knockout mice. These results challenge previous hypotheses about red blood cell, monocyte, or endothelial sources of Arg1 in malaria. *Plasmodium* parasites also encode an arginase that significantly depletes arginine *in vitro* (Olszewski *et al*. 2009) but not plasma arginine *in vivo* (Alkaitis *et al*. 2016). Consistent with these studies, our results suggest that the host, rather than the parasite, is the source of plasma arginase and arginine depletion in malaria. This, along with the relatively few strong parasite-metabolite correlations we observed, suggests the host is more directly responsible than the parasite for driving plasma metabolic changes in acute malaria.

Many studies observe decreased plasma arginine during infection (Yeo *et al*. 2007, Yeo *et al*. 2009, Yeo *et al*. 2013, Yeo *et al*. 2015, Miller *et al*. 2013, Chau *et al*. 2013, Alkaitis *et al*. 2016, Weinberg *et al*. 2016, Cordy *et al*. 2019, Leopold *et al*. 2019a, Leopold *et al*. 2019b, McDonald *et al*. 2018) and even attempt to restore levels of this vasoprotective amino acid (Yeo *et al*. 2007, Yeo *et al*. 2013, Rubach *et al*. 2019), but the cause(s) of arginine depletion in malaria are still debated. Anorexia and urea cycle changes accounted for some but not all infection-induced plasma arginine depletion in mice with experimental cerebral malaria (Alkaitis *et al*. 2016). Our data support a minor role for anorexia in depleting plasma arginine, and suggest that disrupting the urea cycle by hepatic Arg1 knockout does not deplete arginine but rather increases it. Instead, our data suggest that even small amounts of circulating hepatic Arg1 can dramatically deplete plasma arginine in diverse mice. Thus, arginase may be relevant even at the low concentrations seen in the *P. berghei* model (Haque *et al*. 2011, Scaccabarozzi *et al*. 2018) and in our *S.* Typhimurium-infected mice (**Fig. S4b**).

We did not identify any single factor that is capable of maintaining normal arginine levels in malarial mice; even *Arg1^fl/fl^; TBG-Cre* mice (with significantly reduced plasma arginase activity) eventually suffered from hypoargininemia. Thus, despite our strong correlative evidence for small amounts of circulating hepatic Arg1 depleting arginine, this raises the possibility that there remains an unidentified factor that is the direct cause of arginine depletion. To better delineate the arginine-depleting role of hepatic Arg1 relative to other factors, future studies may benefit from careful kinetic analysis of arginine depletion following arginine infusion in *P. chabaudi*-infected *Arg1^fl/fl^; TBG-Cre* and *Arg1^fl/fl^; TBG-GFP* mice, which theoretically differ only in their levels of circulating plasma arginase.

Our results suggest that liver damage may explain variability in arginine depletion in human malaria patients. Like arginine depletion (**Fig. 5**), liver damage is prevalent but not universal in malaria patients (Reuling *et al*. 2018). Failure to account for differences in liver damage and plasma arginase activity may explain why arginine infusion does not always restore plasma arginine in malaria patients (Yeo *et al*. 2007, Yeo *et al*. 2013). While liver damage and plasma arginase activity did not predict survival across the eight mouse strains used in our study, increased plasma arginase and arginine depletion are associated with pathologies we did not measure, including vascular stress (Yeo *et al*. 2009, Morris *et al*. 2005), negative birth outcomes in pregnant women (McDonald *et al*. 2018), and gut vascular permeability in *Plasmodium*-*Salmonella* co-infection (Chau *et al*. 2013). Given the increased plasma arginase and depleted arginine we observed in *S.* Typhimurium-infected mice, these pathologies may not be limited to malaria. Collectively, our data motivate further study of the relationships between liver damage, arginase, and vascular health in infectious disease. C57BL/6 and CAST/EiJ mice, which had high and low ALT, respectively, may provide a useful model for those studies.

### Future directions

Should liver damage emerge as an important player in hypoargininemia in human malaria, NETosis (Knackstedt *et al*. 2019) and IL-1-mediated inflammation (de Menezes *et al*. 2019) may be therapeutic targets for limiting liver damage. Future efforts might also determine why some non-resilient mouse strains had weak overall cytokine responses while others had hyperinflammatory responses. These data point to a tradeoff between effective immunity and immunopathology in malaria that should be explored. We characterized other strain-specific responses that suggest non-C57BL/6 mouse strains will be the best model for some malarial host responses. We used genetically diverse mice because they displayed high phenotypic diversity, but we did not determine how genetics contributed to phenotypic diversity. Quantitative trait locus mapping in malarial DO mice would achieve that goal. Finally, the insights we gained in this study relied heavily on correlative data that do not link processes across time. To identify metabolic processes that connect early, acute, and late parts of the host response (Lissner *et al*. 2020), longitudinal metabolic sampling of diverse mice would enable use of predictive tools like hidden Markov models.

### Implications and Conclusion

We saw that humans and diverse mice exhibit variation in metabolic responses to malaria that depends on disease severity. This implies that future metabolic studies of infection should account for both the nature and severity of disease. We used diverse mice to highlight the variability of metabolic responses and disease severity following infection, and used this variability to yield new insights about liver damage and arginine metabolism. Collectively, our data provide new bioinformatic and mechanistic insights into infection metabolism and motivate the use of genetically and phenotypically diverse mice in future studies.

## Methods

### 1. Ethics statement

Mouse studies were conducted at Stanford University in accordance with NIH guidelines, the Animal Welfare Act, and federal law. They were approved by the Animal Care and Use Committee. Stanford University is accredited by the International Association for Assessment and Accreditation of Laboratory Animal Care (AALAC). All animal work was done according to protocols approved by Stanford University’s Administrative Panel on Laboratory Animal Care (APLAC) and overseen by the Institutional Animal Care and Use Committee (IACUC) under Protocol IDs 30923 (D.S.S., malaria experiments) and 31047 (D.M.M. *Salmonella* experiments).

### 2. Parasites

*Plasmodium chabaudi chabaudi* AJ (Malaria Research and Reference Resource Center [MR4]) was tested for contaminating pathogens prior to use.

### 3. Mice

Wildtype mice were purchased from the Jackson Laboratory (Bar Harbor, ME, USA) (WSB/EiJ stock #001145), 129S1/SvImJ stock #002448, NZO/HILtJ stock #002105, CAST/EiJ stock #000928, A/J stock #000646, NOD/ ShiLtJ stock #001976, and PWK/PhJ stock #003715), and Diversity Outbred (stock #009376, Churchill *et al*. 2012) Charles River (C57BL/6). A subset of expensive mice (WSB/EiJ, NZO/HILtJ, and PWK/PhJ) were also bred in-house. For *Salmonella* experiments, DO mice (same as *Plasmodium* experiments above) were purchased from JAX. Animals were maintained specific-pathogen free (SPF) and housed in the Stanford Research Animal Facility according to Stanford University guidelines, accredited by the Association of Assessment and Accreditation of Laboratory Animal Care (AAALAC) International. All mouse experiments were approved by the Stanford Administrative Panel on Laboratory Care (APLAC).

AhR knockout mice were purchased from Taconic (C57BL/6-*Ahr^tm1.2Arte^*; stock #9166, B6-F for wildtype controls). Arg2 (*Arg2^tm1Weo^*/J, stock #020286, Shi *et al*. 2001) knockout mice were purchased from JAX. All controls were age-matched and purchased from JAX (JAX B6 stock #000664). Arg1 floxed mice were purchased from JAX (C.B6-*Arg1^tm1Pmu^*/J, stock #008817, El Kasmi *et al*. 2008) and Cre mice (*Tie2:* B6.Cg-Tg(Tek-cre)12Flv/J, stock #004128, Koni *et al*. 2001) were bred in-house. Arg2 mice were further bred in-house to yield littermate control animals.

### 4. *P. chabaudi* infection

2 female C57BL/6 mice were given intraperitoneal (i.p.) injections of 100uL frozen stock of *P. chabaudi*-infected red blood cells (iRBCs). When parasitemia reached 10-20% at 8-10 days post-infection, mice were euthanized and blood obtained via cardiac puncture. Blood was diluted to 10^5^ iRBC / 100uL in Kreb’s saline with glucose (KSG) and administered i.p. to experimental animals at a dose of 10^5^ iRBCs. https://www.nature.com/articles/nprot.2011.313 Control animals received 100uL of vehicle i.p. Only female mice 8-12 weeks of age were used for *P. chabaudi* experiments. Experiments were performed in multiple cohorts. Parasitemia was quantified via thin blood smear, methanol fixation, KaryoMAX Giemsa (GIBCO) staining, and manual microscope counting at 100X magnification. RBCs were quantified using a BD Accuri C6 Plus cytometer (see Longitudinal Infection monitoring).

### 5. *Salmonella* infection

10-week-old females were infected with wild type *Salmonella enterica* serovar Typhimurium SL1344 by oral gavage (1x10^8^ CFU in a volume of 0.1 ml). 28 days after infection mice were humanely euthanized and blood was collected by cardiac puncture in an EDTA coated tube (cat # 22-030-402, Greiner Bio-One North America, Inc., Monroe, NC, USA). For plasma separation, blood was centrifuged at 3000xg 10 minutes at 4C. Samples were kept at -80C until further analysis. All animal experiments were approved by the Stanford University Administrative Panel on Laboratory Animal Care (APLAC) with oversight by the Institutional Animal Care and Use Committee (IACUC).

### 6. Bacterial preparation

Wild type *Salmonella enterica* serovar Typhimurium SL1344 were grown in LB miller (Fisher Scientific) supplemented with 200 μg/mL streptomycin (Gold Bio) for 16 hours with 200rpm agitation at 37C. For mouse inoculation, bacteria were diluted into PBS.

### 7. Longitudinal *P. chabaudi* monitoring

Longitudinal monitoring was performed as described previously (Torres *et al*. 2016). For each mouse, baseline RBC, weight, body temperature, and blood glucose measurements were collected between 1 and 5 days prior to infection. In some cases, blood glucose was collected only at baseline sampling and on the day of sacrifice. Mice were restrained during sample collection using tail-access rodent restrainers (Stoelting Co.). Blood was collected from the tail vein by nicking the end of the tail with disinfected surgical scissors, and depositing the blood into EDTA-coated capillary tubes to prevent clotting. For total RBC quantitation, 2uL of blood was diluted in 1mL of cold 1x Hank’s Balanced Salt Solution (HBSS) and kept on ice until absolute RBC counts were obtained using forward and side scatter gates on a BD Accuri C6 Plus flow cytometer. To record body temperature, mice in the metabolic screen experiments were implanted with subcutaneous electronic temperature and ID transponders (IPTT-300 transponders, Bio Medic Data System, Inc) one week prior to infection. Mice were locally anesthetized using a 2% lidocaine solution (100 ug delivered per dose) prior to implantation. Temperature data was recorded using a DAS-7006/7 s reader (Bio Medic Data System, Inc). Subsequent to metabolic screen experiments, body temperatures were measured using a thermocouple thermometer and mouse rectal probe (World Precision Instruments, RET-3). Blood glucose measurements were obtained with 2uL of tail vein blood analyzed with a Bayer CONTOUR Blood Glucose Monitor and Test Strips. Post-infection sampling began on day 4 or 5 post-infection. Parasitemia values were obtained as detailed above. Parasite density is the number of iRBCs per microliter of blood, and is calculated by multiplying parasitemia by the number of total RBCs.

### 8. Flow cytometry

White blood cells, platelets, and reticulocytes were quantified in a subset of mice at baseline sampling, and on odd-numbered sampling days from day 3 through day 11. Cells were first quantitated on an Accuri cytometer as detailed in “Longitudinal *P. chabaudi* monitoring.” RBC counts were used as an approximation for total blood cell counts. Approximately 10 million cells were plated in FACS buffer (PBS, 0.2% fetal bovine serum (Sigma), 5 mM EDTA). Prior to staining, the cells were incubated in TruStain FcX antibody (Biolegend) for at least 5 min at 4 °C. A cocktail containing the Live/Dead Fixable Blue stain (Fisher L34962) and antibodies against the following antigens was added to the blocked cells: CD71 PerCP-Cy5.5 (clone RI7217), TER-119 PE-Cy7 (TER-119), TCRgd PE (UC7-13D5), CD19 Brilliant Violet (BV) 785 (6D5), CD3 BV650 (17A2), CD8 BV510 (53–6.7), Ly6G BV421 (1A8), CD4 Alexa Fluor 700 (GK1.5), Ly6C Alexa Fluor 647 (HK1.4), CD335 FITC (29A1.4) (all from Biolegend); CD11b Alexa 780 (M1/70, eBioscience); CD41 BUV395 (MWReg30, BD Biosciences). All stains were performed for 12–15 min at 4 °C. 5 ml of CountBright counting beads (Invitrogen) were added to each sample such that absolute counts per ml of blood could be back calculated. Data were acquired on an LSR Fortessa (BD Biosciences) and analyzed using FlowJo (Tree Star). Prior to sample acquisition, splenocytes were obtained from a healthy mouse spleen, stained with *α*-CD4 antibodies, and used for instrument compensation.

### 9. Plasma collection for cross-sectional analyses

Between 3 and 5 infected mice of each strain were euthanized each day from days 3-12 post-infection for cross-sectional analysis. Because the WSB/EiJ strain experiences delayed peak infection severity relative to the other mouse strains in this study, 3 or 4 WSB/EiJ mice were euthanized each day from days 3-17 post-infection for cross-sectional analysis. For each mouse strain, 2 uninfected control animals were euthanized at baseline and generally on odd-numbered days between days 3-12 or 3-17 for WSB/EiJ mice. Euthanasia was performed using carbon dioxide asphyxiation in accordance with Stanford University and APLAC guidelines for humane euthanasia. Following euthanasia, blood was collected via cardiac puncture using 25Gx5/8IN tuberculin syringes (Fisher Scientific 14-841-34). Syringes were primed by filling the syringe barrel with 0.5M EDTA, pH 8.0 anticoagulant and dispensing all but 50uL. Collected blood was stored on ice in 1.5mL Eppendorf tubes for 15-45 minutes before spinning at 1,000xg at 4 degrees C for 5 minutes in a tabletop centrifuge. Plasma was frozen at -80C immediately, and thawed/re-frozen once to aliquot for downstream cytokine, metabolite, and liver enzyme analyses.

### 10. Cytokine analysis

75uL of plasma was sent to the Human Immune Monitoring Center at Stanford University. Mouse 38-plex kits were purchased from eBiosciences/Affymetrix and used according to the manufacturer’s recommendations with modifications as described below. Briefly, beads were added to a 96-well plate and washed in a Biotek ELx405 washer. Samples were added to the plate containing the mixed antibody-linked beads and incubated at room temperature for one hour followed by overnight incubation at 4°C with shaking. Cold and room temperature incubation steps were performed on an orbital shaker at 500-600 rpm. Following the overnight incubation, plates were washed as above and then a biotinylated detection antibody was added for 75 minutes at room temperature with shaking. Plates were washed as above and streptavidin-PE was added. After incubation for 30 minutes at room temperature a wash was performed as above and reading buffer was added to the wells. Each sample was measured as singletons. Plates were read using a Luminex 200 instrument with a lower bound of 50 beads per sample per cytokine. Custom assay control beads by Radix Biosolutions were added to each well. Samples in which >50% of cytokines returned low bead counts (<=25) were filtered from the dataset. Individual datapoints with bead count <=25 were also removed. For each cytokine, raw median fluorescence intensities are reported as the number of standard deviations from the mean of all samples (Z-scores).

### 11. Metabolite analysis

100uL of plasma was shipped to Metabolon (https://www.metabolon.com/, Durham, NC, USA), which performed a combination of gas and liquid chromatography with mass spectrometry (GC/LC-MS). Compounds were identified by comparing sample peaks to an internal Metabolon library of known and unknown compounds. Raw peak values were obtained using area-the-curve. Additional data normalizations were performed by Metabolon to account for sample dilutions and day-to-day variation in instrument performance. The three murine metabolomics experiments in this study (the 8-strain *P. chabaudi* experiment, the *Salmonella* experiment, and the *Ahr^-/-^ P. chabaudi* experiment) were measured in three separate Metabolon sample batches.

### 12. Liver enzyme analysis

In the cross-sectional, 8-mouse strain experiments, 90uL of plasma was sent to the Veterinary Service Center Diagnostic Lab at Stanford University. Parameters measured were plasma markers of liver damage: aspartate (AST) and alanine (ALT) transaminases. Units are reported as units per liter (U/L). In follow-up experiments, ALT was measured in 1 microliter of plasma collected via cardiac puncture or tail vein bleed using a colorimetric assay (Millipore Sigma cat. # MAK052) according to the manufacturer’s instructions.

### 13. Comparison of mouse and human metabolic data

We obtained two human metabolomic datasets to compare with our murine data. The first dataset comprised samples from *P. falciparum*-infected pediatric Malawian patients with signs of cerebral malaria (Gupta *et al*. 2017). Control samples were obtained from the same patients during the convalescent phase of infection. Metabolic data were obtained from Metabolon and the scaled imputed ion counts were used for subsequent Z-score transformation and analysis (see below). The second dataset is derived from *P. falciparum*-infected Thai adults in Experiment HuB from the Emory University Malaria Host-Pathogen Interaction Center (MaHPIC) public data releases: http://www.systemsbiology.emory.edu/research/Public%20Data%20Releases/index.html. Only targeted metabolomics data (Biocrates Life Sciences AG, Innsbruck, Austria) were used for this analysis. Murine data were obtained from Metabolon as described above. To compare among datasets, metabolite names were manually matched and only metabolites measured in all 3 datasets (n=42) were included in subsequent analysis. For each metabolite, values were converted to Z-scores by dividing by the mean of values from uninfected C57BL/6 mice (this work), from convalescent individuals (Gupta *et al*. 2017), or from healthy non-malarial control patients (MaHPIC). For mouse samples only, we summarized metabolite Z-scores by reporting the median Z-score for each day and mouse strain. For network analysis, we generated a sample-wise Pearson correlation matrix using all Z-scored *P. falciparum*-infected human samples and median Z-scored mouse samples from acute infection (days 14-17 for WSB/EiJ or days 8-11 for other strains). In the undirected network, each sample is a node, and edges were drawn between nodes whose correlation coefficients were >0.707 (R^2^ > 0.5). We used the same Z-scored metabolite values in the dot plots that show individual metabolites.

### 14. PCA and CCA

Principal component analysis and canonical correlation analysis was performed using the phyloseq R package (McMurdie & Holmes 2013) with Bray-Curtis dissimilarity as the distance metric. Infected and uninfected samples from all days and mouse strains were included in the analysis. Scaled imputed ion counts were Z-scored using uninfected C57BL/6 values as the mean, and a pseudocount was added to all values to make the minimum value = 1 (Bray-Curtis dissimilarity cannot be computed on negative values). Only metabolites present in >= 80% of samples were included in analysis, and bile acids were removed from analysis prior to correlations because many were inconsistently detected across strains. For CCA for both infections, any metabolite with vector length > 0.1 in CA1 or CA2 was included in the comparison to identify metabolites that overlap between infections.

### 15. Correlation analysis

Pearson correlations were performed on log_2_-transformed disease severity metrics and log_2_-transformed scaled imputed ion counts. Bile acids were removed from analysis prior to correlations because many were inconsistently detected across strains. A total of 751 metabolites were included in the final correlation analysis. P-values for correlations were Bonferroni-corrected. All samples (infected and uninfected from all days and mouse strains) were included in correlation analysis.

### 16. sPLS-DA

Partial least squares discriminant analysis (PLS-DA) is a supervised classification tool that performs dimensionality reduction and variable selection. Sparse (s)PLS-DA is a sparsity-penalized version of PLS-DA that is often applied to ‘omics’ data. We used the mixOmics R package (Rohart *et al*. 2017) implementation of sPLS-DA to identify metabolites (“features”) that covaried with ALT values, a proxy for liver damage. We first classified samples as low (ALT <= 100 U/L), medium (ALT 101-1000 U/L), or high (ALT > 1000 U/L) liver damage. As in the correlation analysis, scaled, imputed ion counts were log_2_-transformed. Samples from CAST/EiJ and C57BL/6 mice from days 7-9 post-infection were included in analysis. Samples from the AhR experiment were from days 7-8 post-infection. We pre-filtered metabolites to remove those with zero standard deviation in uninfected animals, and to remove bile acids because many were inconsistently detected across strains. Component selection (n=3 for both datasets) was chosen to minimize error rates. sPLS-DA is run multiple times on different sample subsets; we selected Leave-One-Out cross-validation to generate these subsets. Feature stability is the frequency with which a given feature was selected during cross-validation. We allowed for 50 features to be selected for each component to detect as many stable features as possible. For comparison between sPLS-DA and correlations, we included sPLS-DA features with stability frequency >=0.9 and correlations with R^2^ >= 0.5.

### 17. AAV

Adeno-associated viral particles were purchased from Addgene (Watertown, Massachusetts, USA). TBG-Cre (AAV.TBG.PI.Cre.rBG, item #107787-AAV8, a gift from James M. Wilson) and TBG-GFP (pAAV.TBG.PI.eGFP.WPRE.bGH, item #105535-AAV8, a gift from James M. Wilson) were thawed immediately prior to use and diluted to in sterile, pyrogen-free Dulbecco’s PBS). Virus was injected IP 7 days prior to *P. chabaudi* infection, and monitoring of mouse body weights began one week post-vector injection. Dosing was based on Ballantyne *et al*. 2016: high = 1 x 10^11^ genome copies (gc), medium = 5 x 10^10^ gc, low = 1 x 10^10^ gc.

### 18. Dietary restriction

Infected and uninfected mice were given food and water *ad libitum*. Experimental treatments for uninfected mice were initiated one day after infected mice (e.g. day 4 post-infection for infected mice was day 3 post-infection for uninfected mice). Food intake, water consumption, and body weight were measured in all mice starting 2 days before infection. Dietary restriction began at day 4 post-infection, when each uninfected mouse was given only as much food as its respective infected control consumed the previous day. *Ad libitum* water access was maintained for all mice at all times.

### 19. Quantitative amino acid measurements

**Analytes and internal standards (ISs)** arginine (ARG; IS = 13C6, 15N4 ARG), ornithine (ORN; IS = d7 ORN), citrulline (CIT; IS = 13C6, 15N4 ARG), asymmetric dimethyl arginine (ADMA; IS = d7 ADMA:HCL), spermidine (SPD; IS = d8 SPD), phenylalanine (PHE; IS = d8, 15N PHE), glutamine (GLUT; IS = d5 GLUT), monosodium glutamate (MSG; IS = d5 GLUT), proline (PRO; IS = d8,15N PHE). Standards were obtained from Cambridge Isotope Laboratories (Tewksbury, MA, USA). **Internal standard preparation**: All reference standards were commercially available in powder form. Stock solutions for each standard (10-200mM) were prepared by solubilization in water and diluted further with water:acetonitrile to 200uM working solutions. **Sample preparation**: For plasma amino acid quantitation, 25uL of frozen plasma was sent for analysis. 5 ul internal standard solution was added to 20 ul plasma aliquot followed by vortexing. 75 ul ice cold solution of acetonitrile/0.1% formic acid was added to the sample, followed by vortexing, then centrifugation. Supernatant was transferred to a new vial and analyzed by LC-MS/MS. **Calibration curves**: Individual analyte primary stock solutions were prepared in water (10-200 mM). Intermediate stock solution (200 uM) consisting of all unlabeled analytes: ARG; ORN; CIT; ADMA; SPD; PHE; MSG; GLUT; PRO, was prepared from individual primary stock solutions. This intermediate stock solution was serially diluted with 50x diluted charcoal stripped plasma (CSP) to obtain a series of standard working solutions which were used to generate the calibration curve. Standard working solutions were prepared freshly for sample analysis. Calibration curves were prepared by spiking internal standard solution (10uM) consisting of six labelled compounds (13C6, 15N4 ARG; d7 ORN; d7 ADMA:HCL; d8 SPD; d8, 15N PHE; d5 GLUT). Because of interference due to endogenous metabolites, calibration curves were prepared in diluted CSP to closely match the study samples. A calibration curve was prepared fresh with each set of samples and it ranged from 0.2 nM to 4000 nM. **Instrumentation:** All analyses were carried out by high-resolution LC-MS/MS using an Agilent Poroshell 120 HILIC-Z column (2.7um particle size, 3.0x100mm) on a Quattro Premier triple quadrupole mass spectrometer (Waters) coupled with a 1100 HPLC system (Agilent). HPLC conditions: column was operated at 30 °C at a flow rate of 0.45 mL/ min. Mobile phases consisted of A: 40mM ammonium formic acid (pH 3.2) in water and B: 40 mM ammonium formate (pH 3.2) in 90% acetonitrile. Elution profile: 90% B for 1.5 min, followed by a gradient from 90% to 30% in 8.5 min, then 30–95% in 1 min, and held at 90% B for a total run time of 16 min. Injection volume was 5 ul. **Quantification**: Selected reaction monitoring (SRM) was used for quantification. Analyte mass transitions are shown in **Chapter 3 Methods Table 1**. Dwell time was 7 ms. Quantitative analysis was done with QuanLynx software (Waters). Calibration curves were linear (R > 0.99) over the concentration range using a weighting factor of 1/X where X is the concentration. The back-calculated standard concentrations were ±20% from nominal values.

### 20. Arginase activity assay

Arginase activity assays were performed using a colorimetric assay (Abcam cat. #180877) according to the manufacturer’s instructions with the following modifications: hydrogen peroxide standards were not incubated at 37 degrees; they were prepared immediately prior to addition of reaction mix. Prior to assay, plasma samples (2.5-10uL) were loaded into Amicon 10kDa spin filters (Fisher Scientific cat. #UFC501096). 400 microliters water was added and filters were spun at 15,500xg at 4 degrees C for 10 minutes in a tabletop centrifuge. After discarding flowthrough, this step was repeated for a total of 2 spins. Retentate was resuspended in arginase activity buffer (provided in Abcam cat. #180877) to a final ratio of 2.5 microliters original sample to 100 microliters total volume. Spin-filtered samples were kept on ice and assayed the same day. To control for inter-assay variability, arginase activity values were adjusted so that the negative control sample for each batch (an aliquot from the same uninfected plasma sample) had arginase activity of zero.

### 21. Data availability

a. data for Founder Strains experiments (disease severity, metabolomics, cytokines, flow cytometry): this paper, Dataset S1. Metabolomics data for the Founder Strains is also available at Metabolomics Workbench, doi: 10.21228/M8RX1D, under title “Plasma metabolomics of diverse mouse strains infected with Plasmodium chabaudi”
b. C57BL/6 WT vs. *Ahr^-/-^* data: Lissner *et al*. 2020
c. *Salmonella* metabolomics data: this paper, Dataset S3
d. data for remaining figures, and scripts for all figures/tables, is maintained at: Github, https://github.com/dave1618/strainsMetabspaper/

## Acknowledgments

We thank Dr. Peter Murray for providing Arg1 null primers for genotyping, Karolina Krasinska & Beryl Xia at Stanford University Mass Spectrometry (SUMS) for data and method development. Thanks to Tsukushi Kamiya and to members of the Schneider, Monack, & Sonnenburg Labs for helpful discussions and reagents. This work was supported by the following funding sources: National Science Foundation Graduate Research Fellowship Program (DGE-1656518, N.M.D.), National Institutes of Health T32 AI007328 (N.M.D. and M.M.L.), Defense Advanced Research Projects Agency W911NF-16-0052 (D.S.S., D.M.M.), a Stanford Discovery Grant (D.S.S.), and a SUMS seed grant (N.M.D., D.S.S.). This research was supported by the Division of Intramural Research, National Institute of Allergy and Infectious Diseases, National Institutes of Health, Bethesda, MD, USA. Funders were not involved in study design, data collection, analysis, or interpretation, writing of the manuscript or decision on where to submit for publication.

## Supplemental material legends

**Table S1. Metabolites whose CCA vectors indicate acute infection in malaria and salmonellosis.** Infected and uninfected samples from all days and mouse strains were included in the analysis. Scaled imputed ion counts were Z-scored using uninfected C57BL/6 values as the mean for malaria data and uninfected DOs as the mean for *Salmonella* data. A pseudocount was added to all values to make the minimum value = 1 (Bray-Curtis dissimilarity cannot be computed on negative values). Only metabolites present in >= 80% of samples were included in analysis, and bile acids were removed from analysis prior to correlations because many were inconsistently detected across strains. For CCA for both infections, any metabolite with vector length > 0.1 in CA1 or CA2 was included in the comparison to identify metabolites that discriminate between health and disease in each infection.

**Table S2. Student’s t-test results.** T-tests were computed to compare the mean values of metabolites in healthy mice from mouse strains that are resilient (WSB/EiJ, NZO/HILtJ, 129S1/SvImJ, C57BL/6) and non-resilient (PWK/PhJ, A/J, NOD/ShiLtJ, CAST/EiJ) to *Plasmodium chabaudi*. Metabolites that differ significantly (p < 0.05 with Bonferroni correction) between resilient and non-resilient strains are shown.

**Table S3. Metabolites identified by sPLS-DA.** Metabolites that stably separate CAST from C57BL/6 samples or *Ahr^+/+^* C57BL/6 from *Ahr^-/-^* C57BL/6 samples on the basis of liver damage using sPLS-DA. “Freq” refers to the frequency with which subsamples of data select a given metabolite during leave-one-out cross-validation (1.00 = 100% of the time). Only metabolites with Freq >= 0.9 are shown.

**Figure S1. Immune responses to malaria vary widely across eight genetically diverse mouse strains. A)** Values of 38 hierarchically clustered cytokines (rows). Each column shows the median values for each of 8 *P. chabaudi*-infected mouse strains on each day of infection (days 0, 3-9 for A/J, CAST/EiJ, NOD/ShiLtJ, PWK/PhJ, days 0, 3-11 for NZO/HILtJ, days 0, 3-12 for C57BL/6 and 129S1/SvImJ, and days 0, 3-17 for WSB/EiJ). Each block of columns corresponds to mouse strain, shown from least resilient (left) to most resilient (right). Within each column block, cells are ordered by day of infection (purple to green, white = day 9). Cell color reflects the median of Z-scored, raw median fluorescence intensity. Gray cells indicate missing data. **B)** Whole-blood flow cytometry for 12 cell types are shown for 7 of 8 Founder Strains. Values for WSB/EiJ were omitted because the TER119 (erythroid lineage) marker failed to mark WSB/EiJ cells. n=3-4 mice per strain per day. The mean of absolute cell counts (+/- standard error) are shown. Note that counting beads were not added to A/J samples; bead counts were estimated using the mean of bead counts from the experiment performed immediately after the A/J experiment.

**Figure S2. Acylcarnitines are elevated in diverse disease states.** 3 datasets were analyzed in addition to our own: LPS-injected female C57BL/6 mice (‘LPS’, from Ganeshan *et al*. 2019), male Wistar rats with acute kidney injury (‘AKI’, from Andrianova et al. 2020), and female C57BL/6 mice under fasting conditions (‘Starvation’, from Pietrocola et al. 2017). These were compared to *P. chabaudi*-infected C57BL/6 (‘BL6’) and CAST/EiJ (‘CAST’) mice from our study on day 9 post-infection. Log2 fold change in metabolite concentration (relative to healthy, uninfected or untreated mice from the source dataset) was calculated for each metabolite. Gray cells indicate missing data.

**Figure S3. Arginase, arginine, and diet dynamics in arginase knockout mice.** (A,B) Mice were injected with increasing doses of TBG-Cre (*Arg1^fl/fl^; TBG-Cre* mice) or a high dose of TBG-GFP (*Arg1^fl/fl^; TBG-GFP* mice) (n=2-3 per treatment). **(A)** Levels of arginine after AAV injection (day - 7) in uninfected mice. Day 0 corresponds to infection day for infected mice in (B). (B) Arginase activity on day 8 post-*P. chabaudi* infection (n=3 per group. 1 mouse in the high- and medium-dose treatments died prior to day 8). **(C)** Plasma arginase activity (nmol H_2_O_2_ / min. / uL blood) and arginine for infected mice on days 7-9 post-infection (1 representative experiment for *Arg1^fl/fl^; Tek-Cre* and *Arg1^fl/fl^* mice, and 2 experimental replicates for *Arg2^+/+^* and *Arg2^-/-^* mice. **(D,E)** Male and female mice (n=3 uninfected and n=4 infected) were injected with high-dose TBG-Cre (red) or TBG-GFP (black). Food intake **(D)** and body weight **(E)** were measured. Each uninfected mouse under dietary restriction (diet restr.) was age-, sex-, and genotype-matched to a *P. chabaudi*-infected mouse and was allowed to eat only as much as its corresponding infected mouse. Infected mice and a subset of uninfected mice were fed *ad libitum* (reg. diet).

**Figure S4. *Salmonella* Typhimurium alters plasma metabolomes and increases plasma arginase activity. (A)** Bray-Curtis dissimilarity index was computed on 623 metabolites per sample for infected and uninfected samples (n=136) from Diversity Outbred (DO) mice. Principal component analysis (PCA) was performed, and sample positions in the first two principal components are displayed. Point colors reflect pre-infection (day 0, pink) values, acute infection (days 3-14, green), and convalescence (days 21-31, blue). **(B)** Plasma arginase activity was correlated with scaled imputed ion counts of arginine (log10-transformed Z-scores) in Diversity Outbred (DO) mice (n=22) on day 28 post-infection with *S.* Typhimurium.

**Dataset S1. Founder Strains malaria experiments.** This file contains data from the Founder Strains experiments, including disease severity, metabolomics and metabolite metadata, and cytokine measurements. It also contains sample metadata, e.g. mouse identifiers and characteristics, and experimental notes.

**Dataset S2. Correlations between disease severity and metabolites.** This file contains the correlation information for each unique combination of disease severity metrics and metabolites shown in Figure 3. It contains R, R^2^, and P values, as well as metabolite metadata.

**Dataset S3. *Salmonella* DO experiments.** This file contains data from the *Salmonella*-infected Diversity Outbred mice, including metabolomics and metabolite metadata, as well as mouse metadata, e.g. identifiers and disease severity.

